# Interplay between resource dynamics, network structure and spatial propagation of transient explosive synchronization in an adaptively coupled mouse brain network model

**DOI:** 10.1101/2023.11.11.566570

**Authors:** Avinash Ranjan, Saurabh R. Gandhi

## Abstract

Generalized epileptic attacks, which exhibit widespread disruption of brain activity, are characterized by recurrent, spontaneous and synchronized bursts of neural activity that self-initiate and self-terminate through critical transitions. Here we utilize the general framework of explosive synchronization (ES) from complex systems science to study the role of network structure and resource dynamics in the generation and propagation of seizures. We show that a combination of resource constraint and adaptive coupling in a Kuramoto network oscillator model can reliably generate seizure-like synchronization activity across different network topologies, including a biologically derived mesoscale mouse brain network. The model, coupled with a novel algorithm for tracking seizure propagation, provides mechanistic insight into the dynamics of transition to the synchronized state and its dependence on resources; and identifies key brain areas that may be involved in the initiation and spatial propagation of the seizure. The model, though minimal, efficiently recapitulates several experimental and theoretical predictions from more complex models, and makes novel experimentally testable predictions.

**Significance statement / Author Summary:** Understanding seizure dynamics at the whole-brain level is crucial for controlling abnormal hypersynchronous activity. Currently, complete brain coverage recordings are lacking in both patients and animal models. We employ network science tools to investigate epileptic seizure-like synchronization in a mouse whole brain network, leveraging network structure and supported dynamics as the basis for seizure evolution. Our results align with experimental findings, suggesting that seizure activity initiates in the cortico-thalamic circuit. Importantly, our novel analysis identifies key nodes, primarily in the cortex, driving this hypersynchronous activity. Our findings highlight network structure’s role in shaping seizure dynamics and the techniques developed here could enhance our control of generalized seizures when combined with patient-specific data.

## Introduction

Epileptic seizures, characterized by bursts of excessive neuronal synchronization which usually self-initiate and self-terminate, are considered as a dynamical disease of brain networks ^1^. Seizures can be classified into distinct subtypes, broadly including those that are confined to a circumscribed area (focal) and those which involve larger sections of the brain (generalized) ^2^. A wide range of microscopic mechanisms contribute to this limited repertoire of seizure types ^3^. Although seizures can originate from different brain regions in patients, they may still manifest similar macroscopic features, as seen in EEG recordings^3–5^. Furthermore, studies have suggested that seizures with similar microscopic mechanisms can present as either focal or generalized depending on the macroscopic network structure ^6^. This decoupling between microscopic and macroscopic dynamics underscores the importance of directly modeling emergent properties of seizures and highlights the significance of adopting a network level approach to studying epilepsy, which has also been recognized by the International League Against Epilepsy ^7^.

Experimental evidence shows that network structure alone is not sufficient but the dynamics supported by it also plays an important role in seizure generation and propagation in a brain network ^8,9^. Seizures have been hypothesized to exist in the bistable regime of dynamical networks that exhibits multiple stable states – normal (unsynchronized) and abnormal (hyper-synchronized). In such a system, random fluctuations (noise) or resource availability can transition the network between the different states, giving rise to transient hyper-synchronized activity seen during epileptic seizures ^1,10^.

The phenomenon of explosive synchronization (**ES**) that is widely studied in complex systems and network science can provide a general framework to understand the role of network structure in facilitating seizure dynamics ^11–14^. ES is characterized by first order, discontinuous and irreversible transitions between globally coherent and incoherent states. These features are highly relevant to seizure dynamics, which also show signatures of critical transitions at both onset and termination across multiple spatial scales ^15,16^. Consequently, ES models have been employed to study abrupt transitions in brain networks ^17,18^. Moreover, complete brain coverage recordings, which can elucidate the dynamics of generalized seizures, are lacking in both patient and animal models. Thus, integrating ES with biological networks enables the study of seizure-like synchronization dynamics at the whole-brain level.

Although ES in complex networks has been successfully modeled using two common microscopic mechanisms – the presence of microscopic correlation features, such as frequency-degree coupling (**FDC**); and adaptive coupling ^12,14^ – they do not explain the transient and recurrent nature of seizures. Experimental and computational studies have linked this transient nature of seizures with the dynamics of energy metabolism ^4,10^. Consistent with this observation, a recent study has shown the occurrence of transient ES (**tES**) in a resource-constrained scale-free network with FDC ^19^, where the time-varying nature of resource consumption is shown to cause the transient behavior.

While resource constrained networks with FDC exhibit tES for certain scale-free networks, there are several other frequently occurring families of network structures (Fig. 1a-c) across which this mechanism does not appear to generalize. Especially in the context of neural dynamics, the network structure of brains (Fig. 1d) is often found to show characteristics of small-world networks (SWNs) as well as scale-free networks (SFNs) (Fig. 1e,f). Moreover, adaptive coupling schemes have been shown to exhibit ES more generally across several network topologies, and in fact ES has been suggested to be a generic property of networks with adaptive coupling ^14^. Adaptive coupling is biologically plausible and has often been used to model interaction in biological systems ^20^. Therefore in this study, we combine resource-constraint with adaptive coupling in a model (Fig. 1g) that can manifest tES across several types of network structures, including classic small-world and scale-free networks as well as a real biological network (a mesoscale mouse brain network, MBN, obtained from the Allen Institute public dataset ^21^).

**Figure 1:**
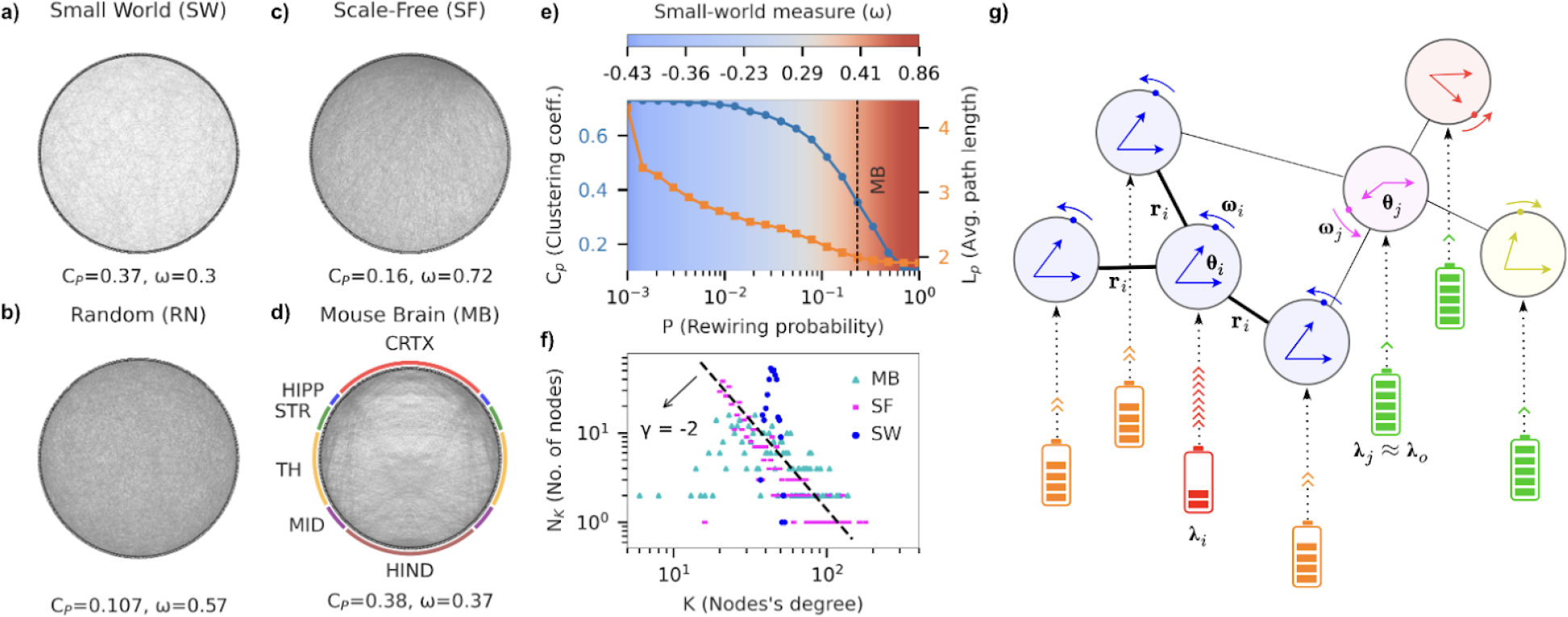
A model with adaptive coupling and resource constraint manifests transient explosive synchronization (tES) in the mesoscale mouse brain network (MBN), that shows both scale-free (SFN) and small-world (SWN) properties. **a**) Visualization of a small-world (SW) network generated using the Watts-Strogatz algorithm (number of nodes, N=400, average degree, ⟨ k ⟩ =40, rewiring probability, *p*=0.232). The generated network has average path length (L_p_ =1.97) and clustering coefficient (C_p_=0.37) similar to mouse brain (MB) network; and small-world measure, ω =0.3. **b)** Random network with path length and average degree matched to MB (L_p_=1.89. ⟨ k ⟩ = 45). **c)** Whole mouse brain mesoscale network having 426 nodes (213 in each hemisphere), each representing a region in mesoscale connectome from Allen mouse brain atlas. The color coded ring around the MB network groups the 426 nodes into 6 major regions (Cortex (CRTX), Hippocampus (HIPP), Striatum (STR), Thalamus and Hypothalamus (TH), Midbrain (MID), Hindbrain (HIND)). The graph (L_p_=2.1, ω =O,37) is generated using a binarized version of the weighted network to allow for comparison with SW and scale-free (SF) networks, **d)** Scale-free (L_p_=1.96) network generated using Barabasi-Alberl algorithm with preferential attachment parameter, m = 20. **e)** Average path length and clustering coefficient for SW network as a function of rewiring probability *p*. The small-world measure ranges between -1 (fully ordered network, blue) to 1 (fully random network, red), with values close to zero (white) corresponding to a perfect small-world network. The clustering coefficient and average path length for MB correspond to a small-world measure of 0.37 (vertical black line), close to 0. f) Degree distribution of SW, SF, and MB networks. Both MB and SF network degree distributions fit the power law distribution with exponent y=-2. SW shows a Gaussiar degree distribution, **g)** (Top) Network of interconnected Kuramoto oscillators, where each oscillator is coupled with every other oscillator through adaptive coupling (a_i_). (Bottom) Resource constraint implies that each oscillator is connected to individual resource reserves (battery), which define the excitability resources of the system (λ_i_). The local synchrony determines the rate of energy consumption as well as the strength of local interactions.

Depending on the resource availability, our model exhibits desynchronized activity, bistability of desynchronized and hyper-synchronized activity (i.e. tES) as well as steady-state hyper-synchronized activity in SFNs, SWNs and the MBN. Furthermore, during the sudden transition to the synchronized state, we observe a wave-like propagation of synchronization across subnetworks within the MBN, beginning with cortico-thalamic subnetworks, followed by subcortical and deeper subnetworks. We also develop a novel algorithm to analyze how the synchronization propagates across individual nodes (brain areas) in the MBN and identify key brain areas that may be responsible for initiation, sustenance and propagation of the hyper-synchronized state. Our results agree with several observations from experimental studies, suggesting that a few key parameters can successfully capture the network level phenomenology of seizure dynamics. Finally the model allows us to study the relationship between the hyper-synchronized state and the resource consumption to recovery rate ratio. Specifically, the model predicts an optimal intermediate ratio for which the likelihood of tES, i.e. epileptic attacks, is minimal. This and related predictions of our model should be directly testable in experiments.

## Results

### An oscillator network model for transient explosive synchronization (tES) based on adaptive coupling and resource constraint

Our model consists of *N* sinusoidal oscillators that form the nodes of a network. Following the Kuramoto model, the interactions between connected oscillators depend on their phase difference. The interaction strength is determined by the structural weight of the connection, and is further modulated by both the synchronization levels of neighboring nodes and the availability of excitability resources (Fig. 1g).

The dynamics of the network are governed by the following equations:

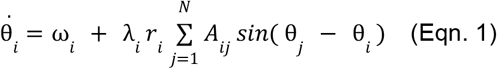

Here, *i ∈* [1, *N*], θ and 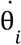 are instantaneous phase and angular velocity of the *i*^*th*^ oscillator, and *ω*_*i*_ is its natural frequency, uniformly distributed in [− 1, 1]. The adjacency matrix *A*_*ij*_ encodes the network structure.

The interaction strengths are modulated by the local synchrony parameter,

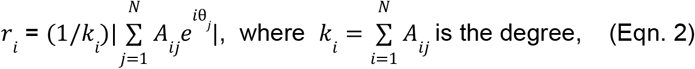

giving rise to adaptive coupling, whereby nodes with higher local synchrony get coupled more strongly.

The interactions are also modulated by the availability of resources to individual nodes, *λ*_*i*_ . Following the model by Frolov & Hramov^19^, we model the time-varying nature of excitability through diffusive coupling as follows:

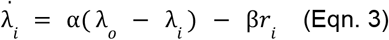

where the first term represents the recovery of excitability resources at a rate α, and the second term represents the local synchrony-dependent resource consumption at a rate β*r*_*i*_ . β is the maximal consumption rate (when *r*_*i*_ = 1). The capacity of the resource reserve for each node is denoted by *λ* (size of resource bath).

The macroscopic behavior of the network is characterized by the global synchrony parameter,

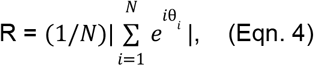

which ranges from 0 (complete desynchronization) to 1 (complete synchronization).

### A Small World Network (SWN) shows tES with adaptive coupling and resource constraint

We begin by investigating the properties of our model in an SWN. For this, we generate an SWN comprising 400 nodes while maintaining parameters such as average degree (⟨k⟩=40), average path length (L_p_=1.97), and clustering coefficient (C_p_=0.37) similar to the MBN for later comparison (Fig. 1a).

We first characterize the resource-dependence of the network dynamics in the absence of resource dynamics (Fig. 2a). Thus, in equation 1, *λ* _*i*_ is replaced by *Λ*, a fixed resource available to each node at all times. We simulate this model with varying *Λ* and observe the steady-state behavior. For this, we begin with *Λ* = 0, adiabatically increase (decrease) *Λ* with increment (decrement) of *ΔΛ* = 0. 003, simulate the model for 1000 time steps and compute the stationary value of global synchrony (R) (see methods), going up to *Λ* = 0.12. For very small *Λ* the network exhibits normal activity, and for very high values of *Λ*, it goes into the hypersynchronized state.

**Figure 2:**
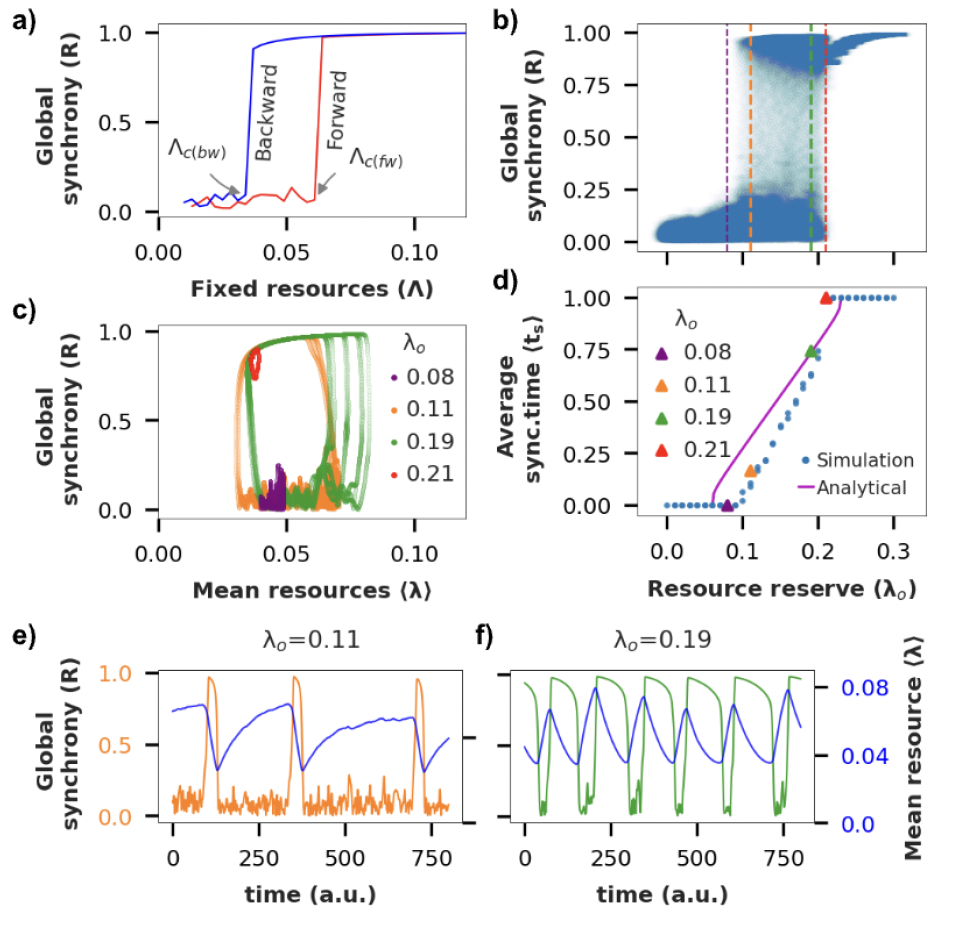
Resource-constrained SWN exhibits tES. **a**) Forward (red) and backward (blue) transition curves for an SWN with adiabatically increasing (decreasing) fixed resource level, Λ Arrows indicate critical points corresponding to forward (or backward) transition from unordered (ordered) to ordered (unordered) state of the network, **b)** Bifurcation diagram of global synchrony (R) vs λ_o_ for SWN with resource constraint, **c)** (⟨ λ ⟩,R) state space trajectory of activity for different values of λ_o_. d) Fraction of time spent in the synchronized state (average synchronized time) vs λ_o_. **e**, **f)** Global synchrony and instantaneous mean resource level as a function of time for two different values of λ_o_ chosen from the bistable region in **(b)**. Note: λ_o_ (fixed) is the fixed resources and < λ > (varying) is the instantaneous mean resources (averaged over all nodes) of the system.

Interestingly for intermediate values, as *Λ* is slowly varied, we observe an abrupt first-order irreversible transition, with the presence of a hysteresis region – depending on the direction of change of *Λ*, or in other words, depending on the current state of the dynamics, the network goes into either the normal or the hyper-synchronized state (Fig. 2a).

Such existence of hysteresis has been shown to give rise to tES when resource constraint is imposed (Eqn. 3) ^19^. When sufficient resources become available, the network transitions to the hyper-synchronized state. This transition is characterized by a sharp decrease in available resources due to increased consumption (Fig. 2d,e). Depending on the resource availability, the network spends a finite amount of time in this state before returning to the incoherent state as resources become depleted. The system replenishes its resources while in the incoherent state and transitions back to the hyper-synchronized state once sufficient resources become available.

Towards testing this hypothesized mechanism for tES, we first identify the parameter range for which the full model shows bistability, i.e. the network spends time in two (meta)stable states (Fig. 2b). We impose resource constraint at a consumption rate β (= 0. 002) and again simulate the full model for *λ*_*o*_, the size of the resource bath, varying between 0.01 and 0.3 with an increment of 0.01. For each value of *λ*_*o*_, we simulate the model for 1000 time steps, and observe the range of values taken by the global synchrony parameter over the simulated period. The resulting bifurcation diagram reveals that the SWN exhibits a globally incoherent state for *λ*_*o*_ < 0. 095, where resources are too limited to allow hyper-synchronization; and a hyper-synchronized state for *λ*_*o*_ > 0. 21. For intermediate values of *λ*_*o*_ the network shows a coexistence of both states, reflecting the presence of tES, as seen in the timeseries of the global synchrony parameter (Fig. 2d,e). Consistent with the proposed mechanism, the transitions to and from the hyper-synchronized state occur when the mean resource availability across all nodes is close to the corresponding critical values of *Λ*, as revealed in the state-space trajectory of the system (Fig. 2c).

Since the transition back to the incoherent state occurs due to resource depletion, we expect the network to spend longer time in the hyper-synchronized state as the size of the resource bath, *λ*_*o*_, increases. This prediction is supported in our simulation results (Fig. 2d) as well as with a simple analytical calculation (see supplement).

Finally we note that the phenomenon of ES or tES is not observed with SWN when correlation feature-based connectivity, such as FDC, is used ^19^.

### The mesoscale mouse brain network (MBN) shows partial tES with adaptive coupling

Next we use our model to study tES in a real biological neural network, using the mouse brain mesoscale connectivity data from the Allen Brain Atlas. The mesoscale atlas is constructed by injecting viral vectors to trace axonal projections across pairs of brain regions in mice ^21^. The dataset consists of detailed and accurate connectivity information across 426 brain areas spanning both hemispheres (see Methods) in healthy mice. The resulting network (Fig. 1d) comprises 11,000 directed edges, with weights rescaled between 0 and 1.

We repeat the analyses from SWN on MBN with an added nuance: while we assumed the SWNs to be binary undirected networks, the dataset we use allows us to define the MBN as a weighted directed network. To account for this, we use a modified version of equations 1 and 2 (see supplement). We analyze all three variants of the MBN – binary-undirected, binary-directed, and weighted-directed. The results presented below refer to the most complete weighted-directed variant unless mentioned otherwise, while the results for the other two variants are qualitatively similar (Fig. S4).

We again begin by characterizing the system with fixed resource availability (see Methods for details). Surprisingly, unlike the SWN which showed hysteresis, the MBN shows bistability for intermediate values of fixed resources (*Λ ∈* [2. 58, 2. 65]) (Fig. 3a). Even binary-undirected and binary-directed versions of MBN, show hysteresis but not bistability for fixed resources (Fig. S4a, S4b). It is important to note that when examining a network with fixed resources, the hysteresis region indicates the potential for tES, but does not guarantee it. On the other hand, the presence of the bistability region directly confirms the existence of tES for the weighted-directed MBN even without the need for resource constraint. We further confirm the existence of tES with simulations of the full model that includes resource constraint (Fig. 3b-d, S3). Note that with the FDC model with or without resource constraints we are unable to induce tES in the MBN (Fig. S4f, S4g).

**Figure 3:**
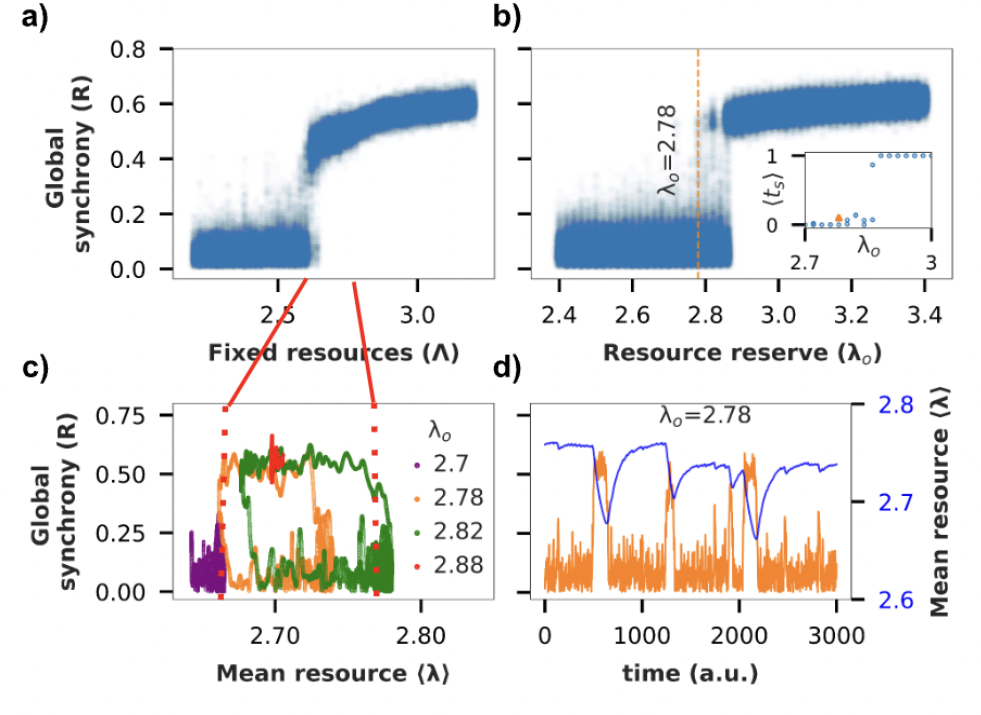
Mesoscale MBN with excitability resource constraint exhibits tES. **a**) Bifurcation diagram of global synchrony (R) vs Λ for the weighted MB network, **b)** Bifurcation diagram of global order (R) vs λ_o_ for the resource constrained model. (Inset) Fraction of time spent in synchronized state vs λ_o_. **C)** (⟨ λ ⟩,R) state space trajectory of activity for different values of λ_o_. **d)** Time series of global synchrony vs instantaneous resources.

Our analysis of the MBN recapitulates the observations from SWN, including the increasing time spent in the synchronized state with increasing size of resource bath (Fig. 3b inset). An interesting deviation is that the synchronized state is only partially synchronized, as reflected in the global synchrony parameter only reaching up to 0.6 instead of 1 (Fig. 3a-d). This happens because certain subnetworks within the MBN never participate in the synchronization. We thus define actively participating nodes as those with a local synchrony greater than the threshold value of 0.7 during the partially synchronized state. We then compare the state space trajectory obtained by averaging the available resources over all nodes versus averaging only over the actively participating nodes (Fig. S2). The state space trajectory shifts to the right in the former case, suggesting that inactive nodes are pushing the average available resources (⟨λ⟩) to higher value. We thus conclude that only the nodes that actively participate in the synchronization cluster increase their energy consumption during tES.

Another deviation from SWNs is that the average resource level (even after restricting to participating nodes) at the time of transitions is observed to be higher than the corresponding resource level in the fixed resource model (Fig. 3a,c). One hypothesis is that even within the synchronized cluster, there may be a core subcluster that drives tES, for which the average energies at transition may be lower, but the peripheral nodes in the synchronized cluster, with higher energy availability, push the apparent average transition energy levels higher. This, however, does not seem to be the case. We conclude that the higher complexity of biological networks, perhaps due to their modular, hierarchical and heterogeneous nature compared to the SWN, makes the relationship between the fixed resource and resource constrained dynamics less predictable.

In the following sections we investigate the dynamics of propagation of synchronization in the MBN.

### ES propagates as a wave from cortico-thalamic to subcortical subnetworks within the MBN

We group the 426 nodes in MBN into distinct communities purely based on the network structure, using the Louvain algorithm (see Methods) ^22^. We hypothesize that such structure-based partitioning of the nodes will group together nodes that are highly likely to form a synchronization cluster. The process naturally partitions the MBN into 8 communities which can be clubbed into three broad classes: cortico-thalamic (4 communities), subcortical (3 communities) and hindbrain (Fig. 4a).

**Figure 4:**
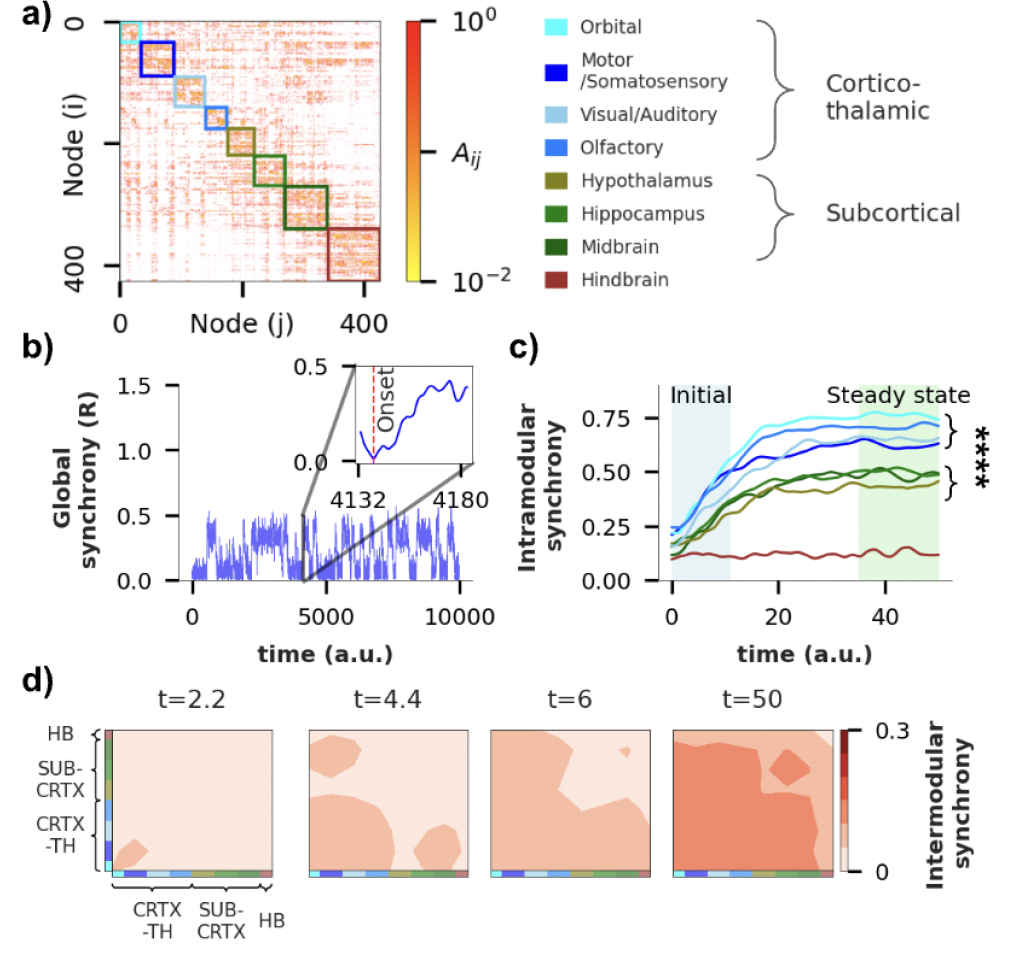
ES propagates as a wave across MBN communities. **a**) Adjacency matrix of MBN where coloured boxes group the 426 nodes into 8 distinct communities obtained using the Louvian algorithm with a resolution parameter of 1. Communities are named based on underlying circuitry (see Supplementary figure 5) and can be further grouped into broader categories: cortico-thalamic (blue) and subcortical (green). Modularity score (Q) of the obtained community partition is 0.53. **b)** The global synchrony parameter reflects several tES events in a weighted MBN for λ_o_ = 2.87. (Inset) activity for one of the transients (out of 52 total) starting from the ‘onset index’ (red vertical line), characterized by the point where complete desynchronization occurs just before the abrupt transition, **c) Intramodular synchrony** (synchronization level within each community averaged over all 52 transients) during the transition shown in (b). Colors represent communities from (a), **d)** Contour map of **intermodular synchrony** for 4 different time points, spanning from a time close to ‘onset index’ (t=2.2) to the time post abrupt transition (t=50) reveals a wave propagating across the network.

We then generate a set of 52 transitions from the incoherent to the hyper-synchronized state by running 4 long simulations, with fixed *λ*_*o*_ = 2. 87, but with different initial conditions (see Methods). Each transition is characterized by the presence of an ‘onset index’, a point where nearly complete desynchronization occurs just before the abrupt transition is about to begin (Fig. 4b, inset), subsequent to which synchronization rapidly increases. To study the spatial propagation of synchronization within the short temporal window in which it occurs, we analyze the transient dynamics over a 50 time unit window following the onset index.

We first study the dynamics of synchronization during the transient window within each community, by computing the intra-modular synchrony (a measure of phase alignment across nodes within the community, see Methods) averaged over all 52 transients. Based on the temporal evolution of the intramodular synchrony, we can group the 8 communities into three distinct cohorts that overlap perfectly with the structural classes defined earlier: cortico-thalamic, subcortical, and hindbrain (Fig. 4c). The steady-state intra-modular synchrony is significantly higher for the cortico-thalamic communities, followed by subcortical, and finally hindbrain, which shows no internal synchronization (*p* < 10 ^−4^ for all the three pairs).

At the start of the transient process, the orbital, olfactory and sensorimotor areas exhibit significantly higher intra-modular synchrony than the rest of the communities (*p* < 0. 05 for each pair, Fig. S8), and also show a steeper rate of increase (Fig. S8), indicating their potential role in initiating and driving the abrupt transition. As the transient progresses, orbital and olfactory areas consistently maintain a significantly higher level of synchronization compared to the rest (*p* < 0. 05, Fig. S8). These results suggest that, while the onset sites of tES can be the orbital, olfactory, or sensorimotor areas, it is likely that the orbital and olfactory areas play a dominant role in driving the transition throughout the entire duration of the transient process. We note that, while intramodular synchrony in the visual area does not seem to significantly differ from the orbital area (Fig. S8), it shows much higher trial to trial variation in terms of its participation. This reinforces previous observations in the literature that the visual area is not critical for the propagation of synchronization^23^.

The dynamics of synchronization propagation can be understood by analyzing the inter-modular synchrony between community pairs (see Methods). Consistent with what we found earlier, ES initiates in the orbital and sensorimotor areas (Fig. 4d, t = 2.2), and then spreads across all cortical regions (Fig. 4d, t = 4.4). Notably, the hippocampal / midbrain areas synchronize with cortical areas before synchronizing among themselves (Fig. 4d, t = 4.4, 6, 50), indicating that the cortical areas are driving their synchronization. At steady state (t = 50), all cortical, thalamic, and subcortical regions achieve synchrony, while hindbrain exhibits minimal participation throughout. Overall, these findings suggest a hierarchical synchronization process, with cortical areas potentially driving synchronization among subcortical regions.

### Propagation of ES across individual nodes in the MBN reveals critical nodes that drive ES

Going beyond the community level, below we assess synchronization propagation at a single node level. Assessing the synchronization between individual nodes typically involves computing the correlation of activity of the pair over short time windows ^24,25^. However, given the extremely short-lived transition window, this technique cannot provide sufficient temporal resolution for studying abrupt transitions. We therefore came up with a novel algorithm that we call the Synchronization Cluster Tracking Algorithm (SCTA) (see methods) for this analysis. The SCTA employs a two-step procedure for quantifying the participation of individual nodes in the synchronization process: (1) generating synchronization clusters by starting from a seed node and expanding the cluster till the local synchrony drops below a fixed threshold; and (2) tracking the synchronization clusters progressively through time to get a ‘cluster lineage’ for each node (see Methods). We apply SCTA to all 52 transients with a local synchrony threshold of 0.5 for the subsequent analysis, although the choice of threshold does not change the results qualitatively (Fig. S5).

The transition typically begins with multiple small synchronization clusters (cluster size < 10) during the early phase (Fig. 5a, t *≈* 17). As the transition progresses, we observe the emergence of a ‘main synchronization cluster’ that quickly comes to dominate in size (Fig. 5a) throughout the rest of the transition. The size of the main cluster varies as the small peripheral clusters or individual nodes continue to join or leave it. This indicates that while the specific onset sites may vary across different trials initially, once the abrupt transition begins, it rapidly spreads from that initial onset site to encompass a broad set of core nodes that hold it together.

**Figure 5:**
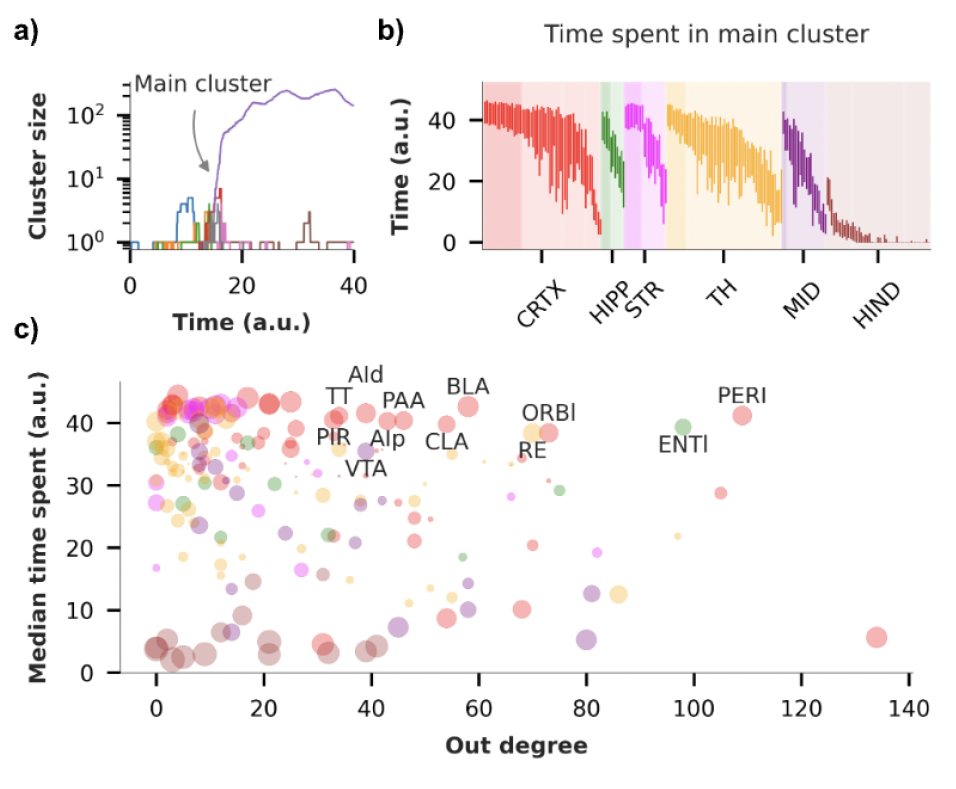
Participation of individual nodes in the main synchronization cluster during the transition. **a**) Synchronization clusters for the transient shown in Fig. 4b obtained using Synchronization Cluster Tracking Algorithm (SCTA) with synchronization threshold of 0.5 (Supplementary material). The figure illustrates one main synchronization cluster (purple, ∼200 nodes) along with several smaller clusters (∼2-20 nodes), **b)** Box plot of the time spent by each node in the main synchronization cluster for 52 different transients (only left hemisphere nodes are shown for clarity). Nodes are color coded as per the major regions defined in Fig. 1c. Dark shaded area in each region shows node with median time spent > 35, and variance in time spent < 49 (σ < 7). **c)** Median time spent in the main synchronization cluster vs out degree of nodes in the left hemisphere (limited to 213 nodes). The size of the scatter points is inversely proportional to the variance in time spent across 52 transients, with larger circles indicating low variance. Labeled points indicate nodes with out-degree > 30, median time spent > 35, and variance in time spent < 49.

To test the ‘core’ nodes hypothesis, we quantify the time spent by individual nodes in the main synchronization cluster across the 52 transients. We find that although a majority of the nodes exhibit participation in the main synchronization cluster, certain nodes consistently spend a significant amount of time in the main cluster (median time spent > 35, std. dev. < 10, Fig. 5b). This core spans across multiple brain areas including the cortex (19 nodes), hippocampus (6), striatum (9), thalamus (10) and the midbrain (3). In contrast, nodes with high variability likely represent peripheral nodes that frequently attach and detach from the main cluster, contributing to the observed variation in cluster size.

This analysis allows us to hypothesize the existence of driver nodes for hyper-synchronization, as nodes with consistent, high participation, and a high out-degree. By spending longer times in the main synchronization cluster, and influencing several downstream nodes they are likely to play a key role in the propagation of synchronization (Fig. 5c). Numerous other nodes with lower out degrees also consistently spend more time in the main cluster, indicating their higher susceptibility to the influence of the drivers, rather than themselves influencing other nodes. These driver areas include the Perirhinal (PERI), Entorhinal (ENTl), Orbital (ORBI), Reticular Nucleus (RE), Basolateral Amygdala (BLA), Piriform (PIR) and Agranular insular area (Ai). Notably, a majority of these “driver nodes” are located in cortical areas. The hindbrain spends the least time in the main cluster, consistent with our earlier findings.

### An intermediate resource recovery-to-consumption ratio is optimal

Since we model resource dynamics explicitly, our model allows us to investigate the impact of resource dynamics on the propensity of tES for the network. In particular, we study the impact of the resource recovery-to-consumption rate ratio (α/β) by fixing the recovery rate (α = 0. 01) and varying the consumption coefficient (β *∈* {0.01, 0.005, 0.002, 0.001, 0.0005, 0.00025, 0.0002}). For this set of parameters, we identify the range of resource bath size (*λ*_*o*_) that support tES. We find that as the recovery-to-consumption rate ratio increases, both SWN and MBN exhibit a shift in the range boundaries to lower values, indicating a higher propensity for tES at low resource levels (Fig. 6a,b). The exponential decrease in range boundaries with an increase in the ratio suggests that even a slight change in the recovery-to-consumption ratio can significantly alter the network’s propensity to generate tES.

**Figure 6:**
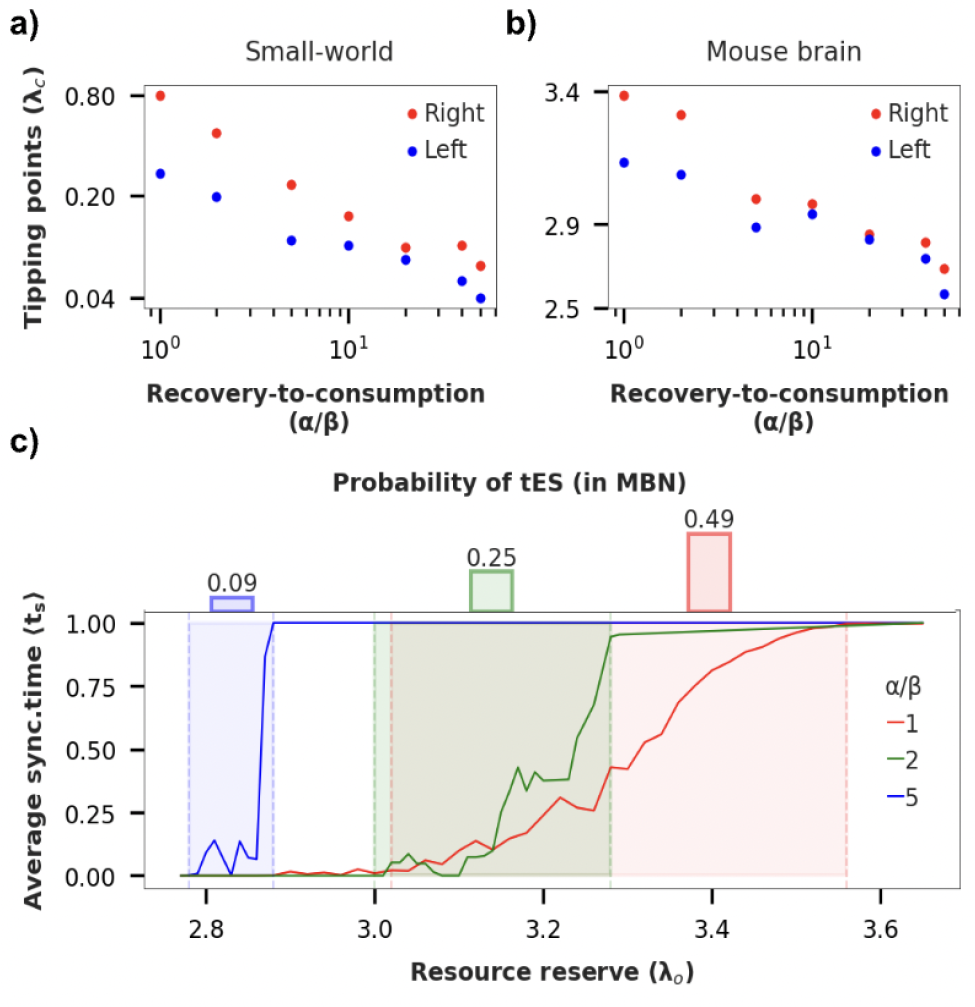
Effect of resource consumption coefficient (β) on network dynamics a, **b)** Critical value of λ_o_ corresponding to left (blue) and right (red) tipping points of the bistability region as function of recovery-to-consumption ratio (or metabolism-to-uptake ratio). Note: Both x and y-axis are on log scale, c) (bottom) Fraction of time spent in synchronized state, (average synchronized time) ⟨t_s_⟩ vs λ_o_ (bistable region shaded), (top) Probability of tES occurrence as function of recovery-to-consumption rate ratio (a/p) (estimated by measuring area under curve in shaded region normalized by area of shaded region).

In contrast, as the recovery-to-consumption rate ratio increases, the width of the bistability region decreases. Moreover, within the bistable region also, the duration of time spent in the synchronized state decreases, resulting in a decreased probability of tES occurrence (Fig. 6c). seizure generation.

Together, these results suggest an optimal intermediate ratio for a healthy brain as follows (Fig. 6d): Assume that the brain always operates at a resource bath size such that it is near the bistable region (criticality hypothesis ^17,26^). Moreover, the resource bath size also undergoes fluctuations. In this scenario, a very low ratio would mean a high probability of tES, i.e. a high likelihood of seizures. If the ratio is very high, the width of the bistable region is so small that fluctuations in the resource bath size push the brain into the monostable hypersynchronized state, again increasing the likelihood of seizures. But an intermediate value of the ratio ensures a balance between these two extremes, and should be observable in a healthy brain.

## Discussion

We first show in an idealized SWN (and other topologies such as SFN) how adaptive coupling can give rise to resource level-dependent hysteresis. Upon the addition of resource dynamics, this gives rise to tES for intermediate sizes of the resource bath.

Recent research comparing diffusive and adaptive coupling, common modeling choices in networks of neural masses, has demonstrated a higher likelihood of networks with adaptive coupling to generate seizures ^20^. Our findings reinforce this preference for adaptive coupling in exhibiting a higher tendency for Our results hold well qualitatively when we apply the same model to a real biological neural network – the mesoscale mouse brain network from the Allen Brain Atlas (Fig. 3). Although the structural network comes from healthy, rather than epileptic mice, our results demonstrate the ability of the model to generate seizure-like dynamics in a biologically realistic network. The framework can be used to further study how perturbations to this network can increase their susceptibility to seizures, thus understanding the specific potential structural elements in diseased mice (or humans) that lead to epilepsy. The choice of a mouse brain as the model system is made because of the greater precision and completeness with which the anatomical connectivity can be measured, compared to non-invasive methods employed in humans.

At the same time, for the MBN we observe some very interesting deviations compared to SWNs. The MBN reaches only a partially synchronized state, with specifically the hindbrain subnetwork never participating in the synchronization (Fig. 3). Even within the synchronized cluster, we hypothesize the presence of a core that becomes fully synchronized, and drives the tES event, and a periphery that does not necessarily reach full synchronization. The energy levels of the nodes at the time of transition may provide a means to identify the core and the periphery. Moreover, the constitution of this core likely depends on the size of the resource bath, so that for different bath sizes, we observe different average transition energies (Fig. 3c). We speculate that this added complexity is a result of the weighted, hierarchical and modular network structure in real brains compared to our idealized SWN. These hypotheses and speculations are areas for further study to understand how the network structure affects its susceptibility to tES.

We then study the dynamics of synchronization propagation across the network at the level of communities (aka subnetworks) and individual nodes. A salient feature we observe is that preceding each abrupt transition, there is a point of near-complete desynchronization across the network (Fig. 4b-inset). This is accompanied by the formation of several small clusters which later merge into the main synchronization cluster. These phenomena are consistent with results obtained through mean field analyses ^14,25^, as well as experimental observations at micro- and macroscopic levels ^27–30^.

At the intra-community level, we find that the cortico-thalamic networks (particularly the orbital, olfactory and sensorimotor areas) exhibit a higher starting synchrony, faster increase of synchrony and a higher steady-state synchrony during the transitions, compared to subcortical communities (Fig. 4c). This suggests their role in initiating and propagating the hyper-synchronized state, consistent with extensive observations and predictions in literature ^31–36^. Additionally, we find certain cortical networks (orbital, sensorimotor) to be more critical for synchronization propagation than others (visual) ^23^.

Quantification of inter-modular synchronization reveals that the synchronization expands in a hierarchical manner, as a propagating wave from the cortical to subcortical regions (Fig. 4d), so that the subcortical areas synchronize with cortical areas before they synchronize among themselves. Although whole-brain recordings during generalized epilepsy are lacking, this would be an interesting hypothesis to test in model organisms with invasive electrophysiology.

We develop a novel algorithm to track the synchronization cluster lineages for individual nodes, which reveals the existence of a single large synchronization cluster during the transition, with several small clusters that dynamically join or leave it. Based on the consistency and time spent by the nodes in the main cluster, and their out degrees, we find a set of ‘core’ nodes that hold the cluster together, irrespective of the initiating site. This driver set includes Perirhinal, Entorhinal, Orbital, Reticular Nucleus, Basolateral Amygdala, Piriform and Agranular insular areas. These predictions are supported by several experimental findings ^37,38^: for instance, the entorhinal, perirhinal, and piriform cortex form a highly interconnected network with other limbic structures and have been shown to possess characteristics that make them susceptible to the initiation and spread of epileptic seizures ^39^.

According to theoretical analysis^14^ and our cluster tracking results, abrupt transitions during tES are preceded by the formation of numerous small synchronization clusters. This is consistently preceded by almost complete desynchronization. The more of these clusters, the more abrupt the transition^14^. A similar phenomenon of synchronization cluster formation and interictal/preictal desynchronization is observed before critical transitions during seizures ^28,29,41^. Experimental evidence shows that these individual clusters exhibit high-frequency oscillations (HFOs) of 80-500 Hz^41^. These observations suggest that the preictal/interictal dynamics of HFO may vary depending on the seizure class that exhibits preictal desynchronization. Testing this hypothesis is intriguing, as it could emphasize the importance of considering seizure type when using HFO as a biomarker _40_.

Lastly, our mechanistic model highlights the importance of an intermediate resource recovery-to-consumption ratio, effectively balancing the heightened tES likelihood and the occurrence of monostable hypersynchronous activity. This implies an optimal recovery-to-consumption range where seizures are infrequent ^42^, and a constant hypersynchronous state is improbable. Deviations from this range may trigger abnormal brain states, suggesting a testable hypothesis for the susceptibility to epileptic attacks in relation to ATP demand and oxygen consumption rates observed during ictal and interictal epileptiform activity ^43,44^.

To summarize, our mesoscale network model for generalized epilepsy applied to a real biological brain network makes several predictions that are consistent with experimental data and more biologically realistic and complex models. The simplicity, coupled with the generality, of the model holds significant value for two key reasons: first, its simplicity allows for the simulation of large-scale brain networks without significant concerns about computational load; and second, it potentially enables the study of seizure dynamics in a wide range of whole-brain networks, and could have applicability from a translational perspective. By identifying the propagation pattern during seizures, we can potentially identify strategies to halt the propagation. Therefore, the model and techniques developed here can be applied to connectome data from actual epileptic brains, with the hope of identifying the seizure onset site and its progression.

## Methods

### Simulations

To evaluate the dynamics of the model with different network structures (small-world, scale-free, mouse brain), we perform two types of simulations: adiabatic progression and bifurcation diagram construction. In the adiabatic progression, we systematically increase or decrease the fixed resource *Λ* to observe the global order of the conventional adaptive coupling model with fixed resources. This allows us to determine the hysteresis region of the system. For the bifurcation diagram construction, we increase the resource bath size parameter *λ*_*o*_ and measure the global order at each time point. For all simulations, we use α = 0. 01, β = 0. 002 (unless otherwise specified) and the initial phases θ*i* are distributed uniformly in the range [0, 2π). Equations are simulated using the Euler method with a step size of 0.05.

To construct hysteresis (bifurcation) diagram in MBN, unlike SWN, we run simulations for a duration of 2000 time units for each value of *Λ* / *λ*_*o*_ through adiabatic progression with *ΔΛ* = 0. 02 / *Δλ*_*o*_ = 0. 02.

To study the progression of ES in a weighted MBN, we conduct four separate runs with a fixed value of *λ*_*o*_ = 2. 87, each using distinct initial conditions for (phase) θ and (frequency) *ω*. The simulations spanned a duration of 20000 time units. From these simulations, we extract a total of 52 transients by selecting segments of length 50-time units, starting from the “onset index” just before the abrupt transition.

## Data Analysis

### Intramodular synchrony

At any particular time unit during the simulation, to asses synchrony level among nodes belonging to one community (obtained from community detection algorithm), intramodular (within community) synchrony, for each community, is computed as the average coherence of phase alignments of all the nodes:

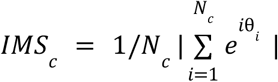

where, *IMS* _*c*_ is intra-modular synchrony is for the *c* ^*th*^ community and *N*_*c*_ is the number of nodes. The average intra-modular synchrony 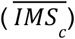 is computed by averaging the synchronization level (*IMS*_*c*_) within each community across 52 transients:

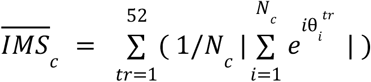

This calculation is performed for each time point within a 50-time unit window to capture the temporal evolution of intra-modular synchrony during the period of abrupt transition.

### Intermodular synchrony

To compute coherence between two distinct communities, intermodular synchrony is computed as the average absolute value of pairwise sum of phase alignment between nodes belonging to different communities:

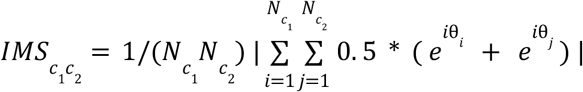

where, 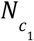 and 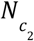 is number of nodes in community *c*_1_ and *c*_2_ . To get temporal evolution of inter-modular synchrony, the average value across 52 transients is computed using a similar methodology as employed for intramodular synchrony analysis.

### Synchronization cluster tracking algorithm

The Synchronization cluster tracking algorithm (SCTA) performs two major tasks: i) finds the synchronization clusters at each time unit (Fig. SM1), ii) tracks the temporal evolution of identified clusters across different time units (Fig. SM2).

The SCTA aims to expand synchronization clusters within the network based on a given synchronization threshold. It follows three key steps:

1. Expansion around Central Nodes: The algorithm begins by expanding the synchronization clusters around the central nodes, which are selected from the previous time step. These central nodes act as trackers for the clusters across different time steps. Nodes with local synchronization exceeding a predetermined threshold become part of the cluster, which terminates with nodes that fall below the threshold (Fig. SM1).
2. Expansion for Unassigned Nodes: Next, the algorithm expands the synchronization clusters for any nodes that have not yet been assigned to a cluster. This step ensures that all nodes are considered and included in appropriate clusters based on the synchronization threshold (Fig. SM2).
3. Update Central Node List: Finally, once all nodes have been traversed or become part of some cluster, the algorithm updates the list of central nodes for each cluster. Central nodes are the top 5 nodes in each cluster with highest local synchrony. These central nodes are important reference points for the clusters, preserving their cluster membership over time, and acting as seed nodes for the expansion of clusters in the next time step (Fig. SM2).

By following these steps iteratively, the algorithm progressively identifies, expands, and tracks synchronization clusters within the network.

**Time spent in main synchronization cluster:**

By executing the SCTA over a 50-time unit window for a specific transient, we obtain the cluster sizes as a function of time. The largest cluster is defined as the main synchronization cluster, and we measure the time spent by each node as part of the main cluster within the 50 time step window.

## Acknowledgements

AKR was supported by a research fellowship from IIT-Delhi. SRG was supported by the Young Faculty Incentive Fellowship from IIT-Delhi. We are also grateful to Dr. Tapan Kumar Gandhi from IIT-Delhi and Dr. Srinivasa Chakravarthy from IIT-Madras for helpful discussions on the topic.

## Author contributions

AKR and SRG conceived the study, designed the model and computational framework, analyzed the data, and wrote the manuscript. AKR carried out the implementation. SRG conceived the idea for the novel algorithm, and in consultation with SRG, AKR further developed and refined the algorithm.

## Code availability

The code and parameters that have provided the results presented here are available at GitHub https://github.com/csndl-iitd/tES_mesoscale_connectivity_model.git

