## Supplementary Material for "Interplay between resource dynamics, network structure and spatial propagation of transient explosive synchronization in an adaptively coupled mouse brain network model"

### Mouse brain network construction and partition

We construct whole brain MBN by combining ipsilateral and contralateral connections from mesoscale atlas of the mouse brain [21], and we select edges with $p < 0.05$. The resulting network comprises 426 nodes and 11,000 directed edges. We normalize the weights of all the edges between 0 and 1 using minmax scaling.

To obtain the modules required for calculating intramodular and intermodular synchrony, we apply the Louvain algorithm [22] to MBN, with a resolution parameter of 1, which groups 426 nodes into 8 distinct communities.

### Modified model for directed network

In the case of directed networks, the local order and adaptive coupling are modified as follows: local order $r_i$ = $1/k_i|\sum_{j=1}^{j\in N_k} e^{i\theta_j}|$, where $N_k$ is all the nodes directly connected to $i^{th}$ node regardless of the direction and $k_i$ is the count of all such nodes in $N_k$. Local interaction via adaptive coupling happens through modified local order $r_i$. These modifications are motivated by the concept of adaptive coupling, which couples nodes exhibiting high local synchrony more strongly. For both weighted and binary networks, the same modified local order is utilized, without considering the weights in the calculation of local order.

For resource-constrained network with frequency-degree correlation features, network is modified as follows:

$$\dot{\theta}_i = \omega_i + \lambda_i r_i \sum_{j=1}^{N} A_{ij} sin(\theta_j - \theta_i)$$

Here, $\omega_i$ is set to $K_i$, where $K_i$ is the sum of in and out degrees for the $i^{th}$ node.

### Average synchronized time

Average synchronized time, $\langle t_s \rangle$, is defined as the fraction of time that network spends in a globally coherent state relative to the total duration i.e. 1000 time units for SWN. To obtain an analytical solution, we utilize the observation that the occurrence of tES in resource-constrained networks is primarily driven by the presence of hysteresis regions (Fig. 2a). Our analysis is confined to a single node, and we employ the prediction from this analysis to compare simulations results obtained at the network level. This comparison is made under the assumption of network homogeneity among all nodes.

#### Time Spent in globally Incoherent State

The time spent in the globally incoherent state is defined as the duration it takes for the variable $\lambda_i(t)$ to transition from point $\lambda_a$ to point $\lambda_b$ within a network operating in a bistable region. Assuming the local synchrony ($r_i$) to be zero during this state results in the following equation:

$$\int_{t_1}^{t_2} dt = \int_{\lambda_a}^{\lambda_b} \frac{1}{\alpha[\lambda_0 - \lambda_i(t)]}$$

, where $t_2$ - $t_1$ is the amount of time network is in incoherent (desynchronized) state ($t_{desync}$). Integrating above equation over the specified limits yields following:

$$t_{desync} = 1/\alpha .\ ln[(\lambda_0 - \lambda_A)/(\lambda_0 - \lambda_B)]$$

**Time Spent in globally coherent State**

Likewise, time spent in coherent state is defined as time it takes $\lambda_i(t)$ to go from $\lambda_b$ to $\lambda_a$ and we assume $r_i$ to be 1 in this scenario. This results in the following equation:

Conversely, the duration spent in a coherent state is defined as the time it takes for $\lambda_i(t)$ to transition from point $\lambda_b$ to point $\lambda_a$. In this scenario, we assume that the rate $r_i$ is equal to 1 resulting in the following equation:

$$\int_{t_1}^{t_2} dt = \int_{\lambda_b}^{\lambda_a} \frac{1}{\alpha[\lambda_0 - \lambda_i(t)] + \beta}$$

, where $t_2$ - $t_1$ is the amount of time network is in incoherent (desynchronized) state ($t_{sync}$). Integrating above equation over the specified limits yields following:

$$t_{sync} = 1/\alpha .\ ln[(\lambda_A - \lambda_0 + \frac{\beta}{\alpha} r_i)/(\lambda_B - \lambda_0 + \frac{\beta}{\alpha} r_i)]$$

**Fraction of time network spends in synchronized state**

To compare the analytical solution with the simulation results, we derive an expression for the proportion of time during which the network resides in a synchronized state relative to the total duration. This is analogous to calculating the average synchronized time (Fig. 2d)

$$\frac{t_{sync}}{t_{sync} + t_{desync}} = \frac{ln\,[\frac{\lambda_B - \lambda_0 + \frac{\beta}{\alpha} r_i}{\lambda_A - \lambda_0 + \frac{\beta}{\alpha} r_i}]}{ln\,[\frac{\lambda_B - \lambda_0 + \frac{\beta}{\alpha} r_i}{\lambda_A - \lambda_0 + \frac{\beta}{\alpha} r_i}] + ln\,[\frac{\lambda_0 - \lambda_A}{\lambda_0 - \lambda_B}]}$$

**Algorithm 1** *Synchronization cluster tracking algorithm*

```
Input:
Synchronization threshold (S_th)
Time window (T_w)
Adjacency matrix (A)
Local order history (L_h)
Output:
node membership history (NM_h)
central_nodes_per_cluster (N_c)← {} : map of cluster id and corresponding 5
  central node list
node_membership_history (NM_h)← []
for t in T_w do
    node_membership ← {}
    filtered_central_node_per_cluster{} ← FilterCentralNodeWithMaxReachabil-
      ity(A, L_h[t], S_th, N_c)
    for node, cluster in filtered_central_node_per_cluster do
        node_membership ← ClusterExpansionAlgorithm(A, L_h[t], S_th, node,
          cluster)
    end
    for node not in node_membership do
        cluster ← GenerateRandomUnusedClusterId()
        NM_h← ClusterExpansionAlgorithm(A, L_h[t], S_th, node, cluster)
    end
    N_c{}← UpdateCentralNodes(L_h, node_membership)
    NM_h[t]← N_c
end
```

**Figure SM1: Synchronization Cluster Tracking Algorithm**

The pseudo code for tracking clusters across the transient process. Initially, clusters are formed using the cluster expansion algorithm. Then, for tracking purposes, five central nodes are selected from each cluster. This selection of multiple central nodes helps to address the issue of detachment of central nodes from the cluster in the subsequent time step. By choosing multiple central nodes, the likelihood of effectively tracking the clusters increases, as even if one of the central nodes remains connected and maintains its membership, the cluster can still be tracked. At the start of each iteration, only one central node with the maximum cluster size is chosen (*FilterCentralNodeWithMaxReachability*).

**Algorithm 2** Synchronization Cluster Expansion Algorithm

**Input:** Adjacency matrix (A)
Local order (L)
Synchronization threshold ($S_{th}$)
Start Node ($N_s$)
Cluster ID ($ID_c$)
Node membership ($N_m$)
**Output:** Node membership ($N_m$)

```
visited nodes (V_n) ← {}
queue (Q) ← deque([start_node])
while Q not empty do
    node ← pop(Q)
    V_n ← add(node)
    if (node not in N_m) and (L[node] > S_th) then
        N_m{node} ← ID_c
        for neighbor_node in A[node] do
            if L[neighbor_node] > S_th then
                if neighbor_node not in N_m then
                    N_m{neighbor_node} ← ID_c
                    Q ← add(neighbor_node)
                end
                else if N_m{neighbor_node} != ID_c then
                    neighbor_cluster_id ← N_m{neighbor_node}
                    N_m{neighbor_node} ← MergeSmallerWithLargerCluster(
                      neighbor_cluster_id, ID_c)
                end
            end
        end
    end
end
```

**Figure SM2: Synchronization Cluster Expansion Algorithm**

The algorithm presented outlines the process of expanding the synchronization cluster at a specific time instance in the MB network. It utilizes a breadth-first traversal (BFT) starting from the start node ($N_s$) and proceeds to expand the cluster in the direction of its neighbors that have a local order greater than the synchronization threshold ($S_{th}$). It iteratively performs the algorithm till all the nodes are visited.

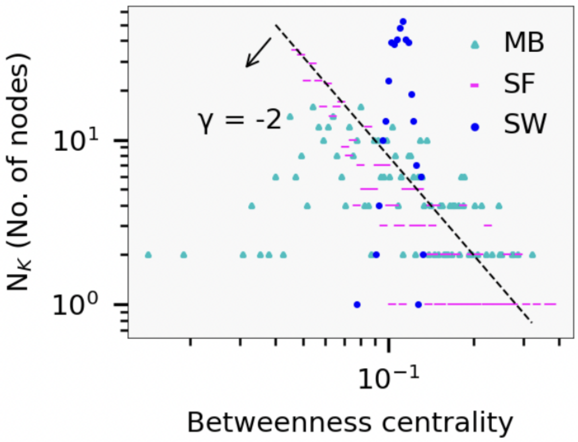


**Figure S1:** Betweenness centrality distribution in MBN, SFN and SWN.

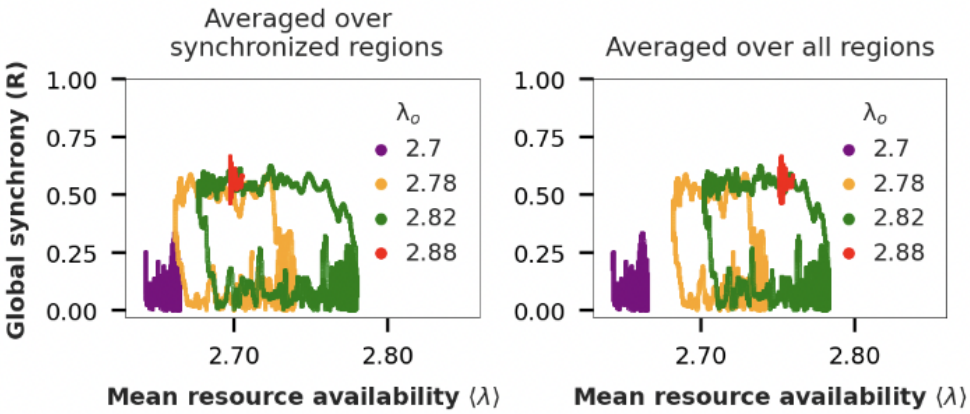


**Figure S2:** (⟨λ⟩,R) state space trajectory of activity for different values of $\lambda_o$. **a)** λ averaged over regions actively participating in transient synchronization events. **b)** λ averaged over all regions.

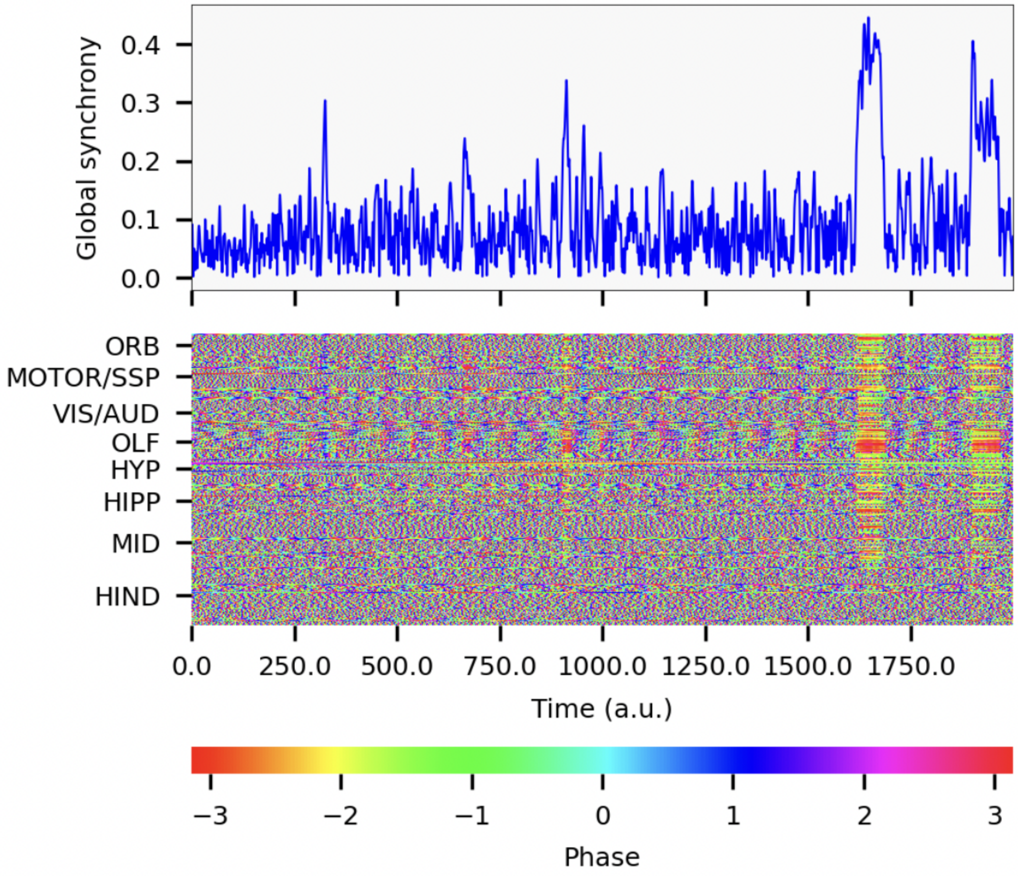


**Figure S3: a)** Time series of global synchrony (R) for fixed $\lambda_o$ (=2.87) in MBN. **b)** Space-time plot of dynamical phases corresponding to **a)**.

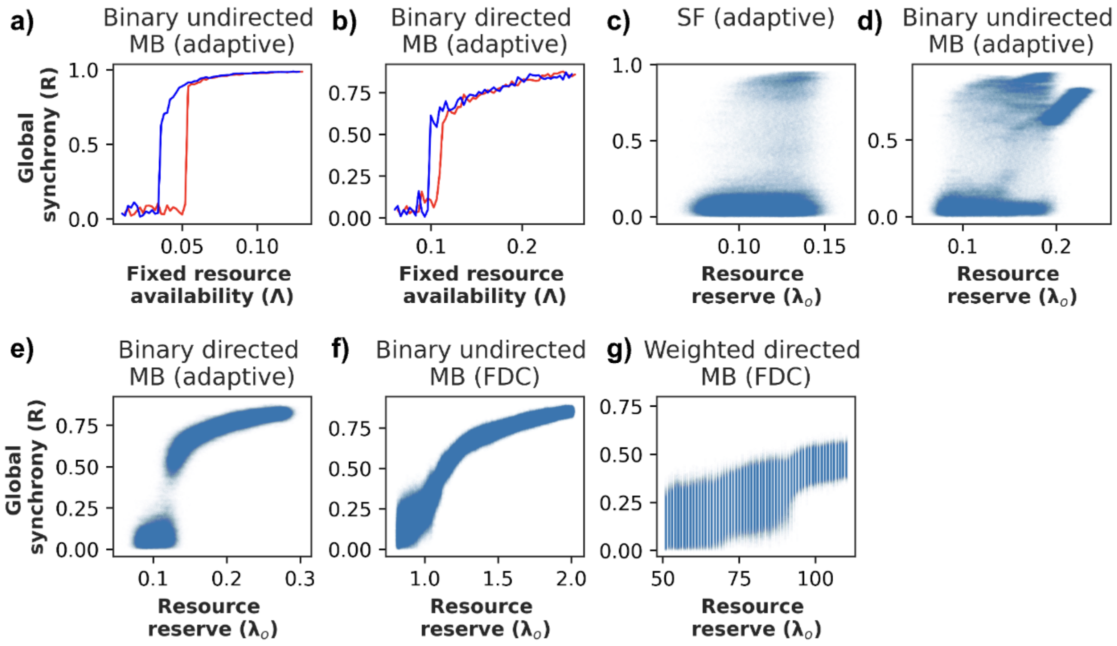


**Figure S4: Hysteresis curve and bifurcation diagram for various network**
Hysteresis curve (**a** - **d**) and bifurcation diagrams (**c** - **g**) for various networks (α=0.01, β=0.002) with adaptive coupling and frequency degree correlation (FDC).

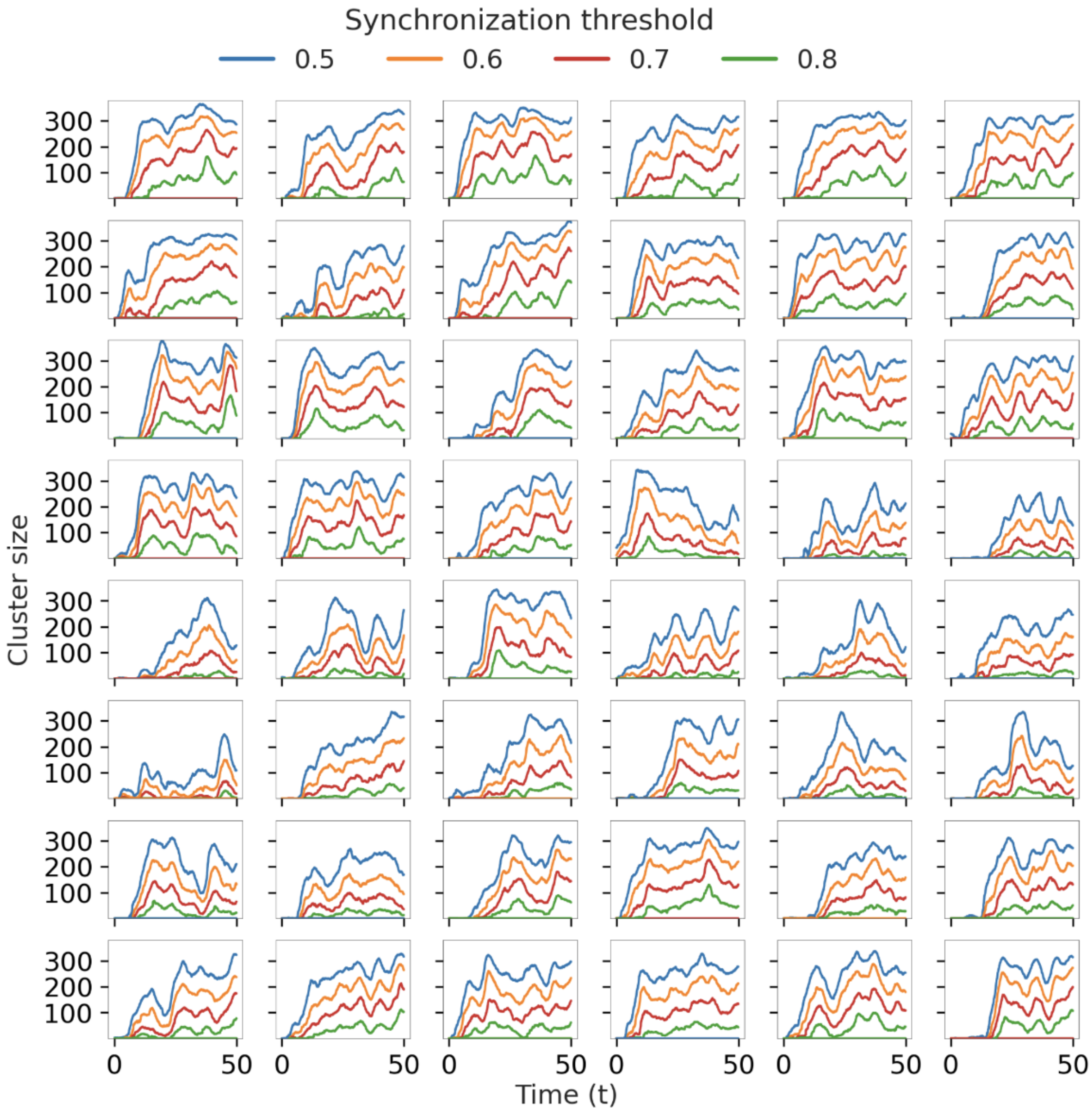


**Figure S5:** synchronization clusters obtained after applying SCTA to 52 different transients (48 shown) for four different synchronization thresholds (0.5, 0.6, 0.7, 0.8).

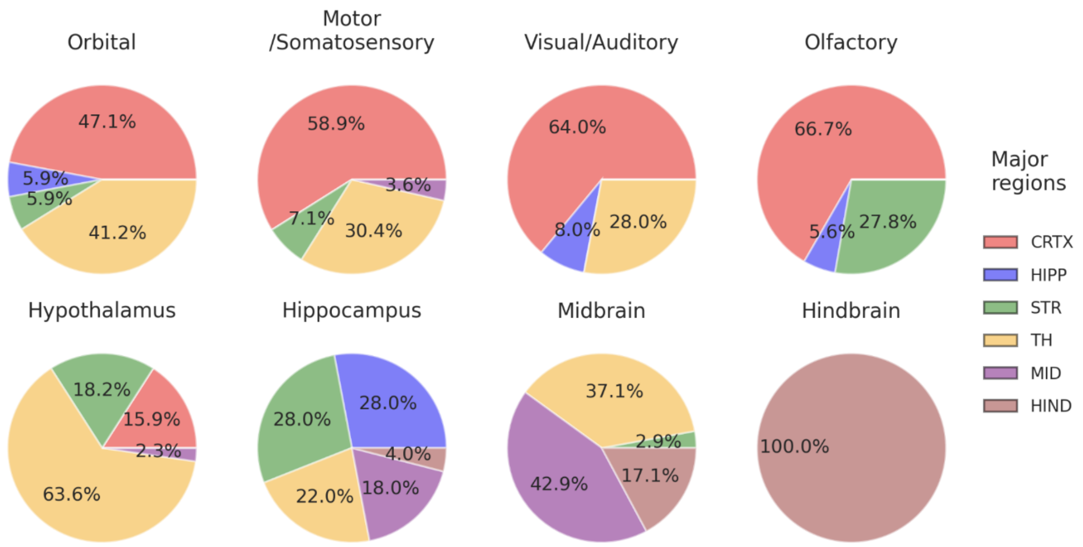


**Figure S6:** Distribution of major regions for each community obtained from Louvian algorithm.

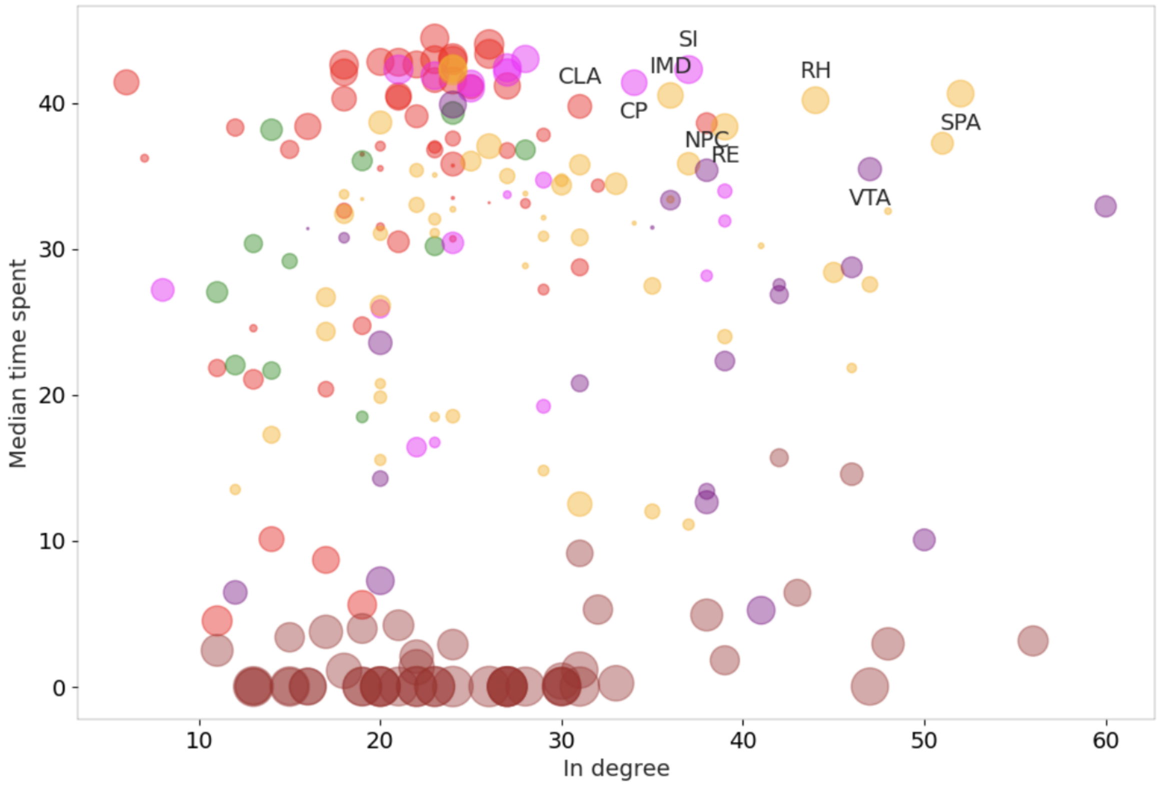


**Figure S7:** The scatter plot displays the relationship between the median time spent in the main synchronization cluster and the degree of nodes in the left hemisphere (limited to 213 nodes). Scatter points are labeled that meet specific criteria: nodes with in-degree > 30, median time spent > 35, and variance in time spent < 49 (σ < 7).

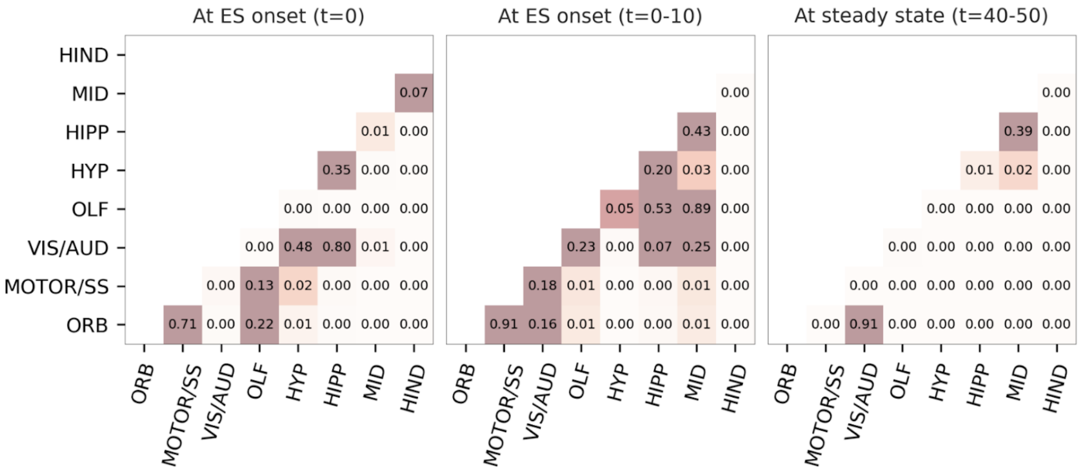


**Figure S8: Weighted t-test on intramodular synchrony at initial and steady state**

P-values obtained by performing **a)** pairwise t-test between intramodular synchrony at tES onset (t=0), weighted t-test distribution for 200 time unit window at **a)** tES onset between slopes of intramodular synchrony, and **b)** steady state between constant steady state values of intermodular synchrony.
